# Streptococcal phosphotransferase system imports unsaturated hyaluronan disaccharide derived from host extracellular matrices

**DOI:** 10.1101/480228

**Authors:** Sayoko Oiki, Yusuke Nakamichi, Yukie Maruyama, Bunzo Mikami, Kousaku Murata, Wataru Hashimoto

**Affiliations:** Laboratory of Basic and Applied Molecular Biotechnology, Division of Food Science and Biotechnology, Graduate School of Agriculture, Kyoto University, Uji, Kyoto 611-0011, Japan; Laboratory of Food Microbiology, Department of Life Science, Faculty of Science and Engineering, Setsunan University, Neyagawa, Osaka 572-8508, Japan; Laboratory of Applied Structural Biology, Division of Applied Life Sciences, Graduate School of Agriculture, Kyoto University, Uji, Kyoto 611-0011, Japan

**Keywords:** glycosaminoglycan, hyaluronan, Streptococcus, sugar import, crystallography

## Abstract

Certain bacterial species target the polysaccharide glycosaminoglycans (GAGs) of animal extracellular matrices for colonization and/or infection. GAGs such as hyaluronan and chondroitin sulfate consist of repeating disaccharide units of uronate and amino sugar residues, and are depolymerized to unsaturated disaccharides by bacterial extracellular or cell-surface polysaccharide lyase. The disaccharides are degraded and metabolized by cytoplasmic enzymes such as unsaturated glucuronyl hydrolase, isomerase, and reductase. The genes encoding these enzymes are assembled to form a GAG genetic cluster. Here, we demonstrate the *Streptococcus agalactiae* phosphotransferase system (PTS) for import of unsaturated hyaluronan disaccharide. *S. agalactiae* NEM316 was found to depolymerize and assimilate hyaluronan, whereas its mutant with a disruption in PTS genes included in the GAG cluster was unable to grow on hyaluronan, while retaining the ability to depolymerize hyaluronan. Using toluene-treated wild-type cells, the PTS import activity of unsaturated hyaluronan disaccharide was significantly higher than that observed in the absence of the substrate. In contrast, the PTS mutant was unable to import unsaturated hyaluronan disaccharide, indicating that the corresponding PTS is the only importer of fragmented hyaluronan, which is suitable for PTS to phosphorylate the substrate at the C-6 position. The three-dimensional structure of streptococcal EIIA, one of the PTS components, was found to contain a Rossman-fold motif by X-ray crystallization. Docking of EIIA with another component EIIB by modeling provided structural insights into the phosphate transfer mechanism. This study is the first to identify the substrate (unsaturated hyaluronan disaccharide) recognized and imported by the streptococcal PTS.

**IMPORTANCE (118/120 words):** The PTS identified in this work imports sulfate group-free unsaturated hyaluronan disaccharide as a result of the phosphorylation of the substrate at the C-6 position. *S. agalactiae* can be indigenous to animal hyaluronan-rich tissues owing to the bacterial molecular system for fragmentation, import, degradation, and metabolism of hyaluronan. Distinct from hyaluronan, most GAGs, which are sulfated at the C-6 position, are unsuitable for PTS due to its inability to phosphorylate the substrate. More recently, we have identified a solute-binding protein-dependent ABC transporter in a pathogenic *Streptobacillus moniliformis* as an importer of sulfated and non-sulfated fragmented GAGs without any substrate modification. Our findings regarding PTS and ABC transporter shed light on bacterial clever colonization/infection system targeting various animal GAGs.

## INTRODUCTION

Extracellular matrices of all animal tissues and organs serve as physical scaffolds for cellular constituents, cell differentiation and proliferation, homeostasis, and tissue formation (1). Glycosaminoglycans (GAGs), constituents of the matrices (2), are acidic polysaccharides consisting of repeating disaccharide units of uronate and amino sugar residues. Hyaluronan, chondroitin sulfate, heparin, and heparan sulfate are classified as GAGs based on their constituent monosaccharides, glycoside linkages, and sulfation patterns (3, 4). Hyaluronan consists of D-glucuronate (GlcUA) and *N*-acetyl-D-glucosamine (GlcNAc), chondroitin sulfates of GlcUA and *N*-acetyl-D-galactosamine (GalNAc), and heparin and heparan sulfate of GlcUA or L-iduronate (IdoUA), and D-glucosamine (GlcN) or GlcNAc (5) (Fig. S1). The uronate and amino sugar residues in hyaluronan and chondroitin sulfate are linked by 1,3-glycoside bonds, whereas the residues in heparin and heparan sulfate are connected by 1,4-glycoside bonds. With the exception of hyaluronan, these GAGs frequently contain sulfate groups in the uronate and/or amino sugar residues, and function as protein-binding proteoglycans in extracellular matrices.

Some bacteria including staphylococci and streptococci target animal GAGs for colonization and/or infection (6). Streptococci are known to invade host cells by the depolymerization of hyaluronan using cell-surface hyaluronate lyase that produces unsaturated disaccharides with C=C double bonds at the nonreducing terminus of the uronate residue, through a β-elimination reaction (7–11) (Fig. 1A). Our previous reports indicate that the resulting unsaturated GAG disaccharides are degraded in the cytoplasm by unsaturated glucuronyl hydrolase (UGL) into monosaccharides (unsaturated uronate and amino sugar), through the hydration of the C=C double bonds (12–14). Moreover, unsaturated uronate was shown to metabolize to pyruvate and glyceraldehyde-3-phosphate through successive reactions catalyzed by isomerase (DhuI), NADH-dependent reductase (DhuD), kinase (KdgK), and aldolase (KdgA) (15). On the other hand, the bacterial import of GAGs is poorly understood, although several ABC exporters of GAGs in bacteria (16, 17) and humans (18) have been identified.

**FIG 1.**
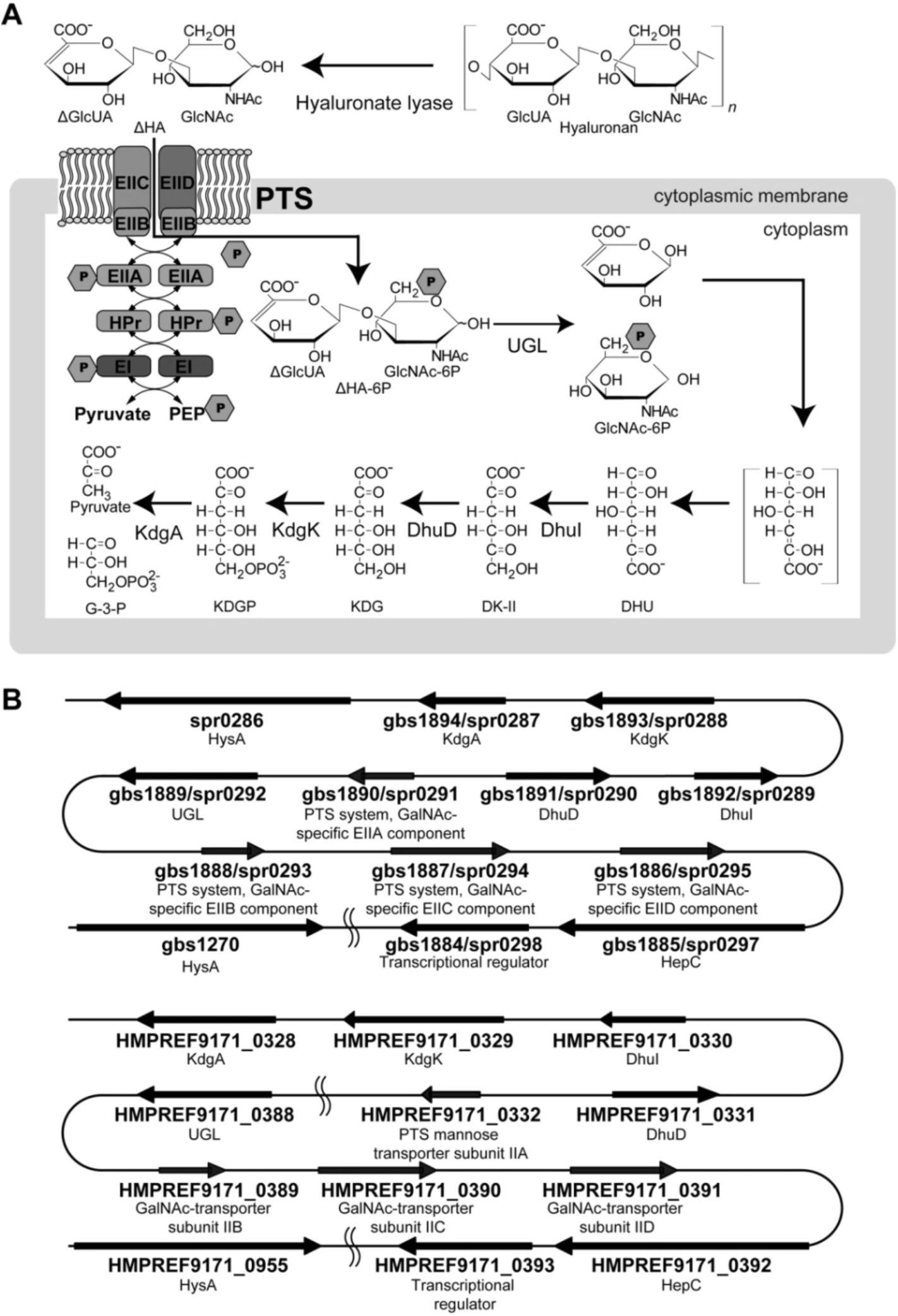
PTS import model and GAG genetic cluster. (A) *S. agalactiae* PTS import model. Cell-surface hyaluronate lyase (spr0286/gbs1270/HMPREF9171_0955) depolymerizes hyaluronan and the resulting unsaturated hyaluronan disaccharides are incorporated into the cytoplasm by the phosphotransferase system (PTS) (spr0291-0293-0294-0295/gbs1886-1887-1888-1890/HMPREF9171_0332-0389-0390-0391). During the import process, a phosphate group is transferred to the substrate. After import has been achieved, unsaturated glucuronyl hydrolase (UGL) (spr0292/gbs1889/HMPREF9171_0388) degrades the disaccharides to monosaccharides. The resulting unsaturated uronate is non-enzymatically converted to 4-deoxy-L-*threo*-5-hexosulose-uronate (DHU). DHU is metabolized to 3-deoxy-D-*glycero*-2,5-hexodiulosonate (DK-II) by 4-deoxy-L-*threo*-5-hexosulose-uronate ketol-isomerase (DhuI) (spr0289/gbs1892/HMPREF9171_0330). DK-II is then metabolized to 2-keto-3-deoxy-D-gluconate (KDG) by 2-keto-3-deoxy-D-gluconate dehydrogenase (DhuD) (spr0290/gbs1891/HMPREF9171_0331). KDG is converted to pyruvate and glyceraldehyde-3-phosphate (G-3-P) via 2-keto-3-deoxy-6-phosphogluconate (KDGP), through successive reactions catalyzed by 2-keto-3-deoxygluconate kinase (KdgK) (spr0288/gbs1893/HMPREF9171_0329) and 2-keto-3-deoxy-6-phosphogluconate aldolase (KdgA) (spr0287/gbs1894/HMPREF9171_0328). (B) GAG genetic clusters in the genomes of *S. pneumoniae* R6 (spr), *S. agalactiae* NEM316 (gbs) (upper), and *S. agalactiae* 5671 (HMPREF9171) (lower).

Streptococci are classified into three groups based on their hemolytic activity: α, incomplete lysis of red cells; β, complete lysis of red cells; and γ, lack of hemolysis (19). β-Streptococci are further classified into A–V groups based on antigenic differences in their cell wall polysaccharides. For example, group α *Streptococcus pneumoniae* is a major causative bacterium of pneumonia, and group β-A *Streptococcus pyogenes* causes pharyngitis and sepsis. Group β-B *Streptococcus agalactiae* is responsible for neonatal sepsis and meningitis, and is indigenous to the gastrointestinal and urogenital tracts of 25–40% of healthy women. In fact, 50% of neonates are maternally infected with *S. agalactiae* during delivery, which can lead to neonatal invasive diseases (20). In *S. pneumoniae* genomes, enzymes for the depolymerization, degradation, and metabolism of GAGs are encoded together with a putative phosphotransferase system (PTS), a typical bacterial sugar import system (21). The genes encoding hyaluronate lyase, UGL, DhuI, DhuD, and the PTS are assembled to form a GAG genetic cluster (Fig. 1B). The similar genetic cluster is also included in the genome of *S. pyogenes* and *S. agalactiae*.

PTS is composed of Enzyme I (EI), histidine-containing phosphocarrier protein (HPr), and Enzyme II (EII), which has multiple hetero-subunits (EIIA, EIIB, EIIC, and EIID) (22). EI and HPr proteins are located in the cytoplasm and nonspecifically recognize sugar substrates, whereas EII is substrate-specific and consists of cell membrane and cytoplasmic domains. Mechanistically, the PTS imports sugar by phosphorylating the substrate at the C-6 position through successive phosphotransfer reactions from a phosphate donor (phosphoenolpyruvate) mediated by EI, HPr, and EII (21). A large number of GAGs (with the exception of hyaluronan) are frequently sulfated at the C-6 position (23). Unsaturated GAG disaccharides with a sulfate group at C-6 are unsuitable as PTS substrates due to the lack of phosphorylation. Indeed, after disruption of the EI gene, *Salmonella typhimurium* still grows on sugars such as glucuronate and glucose-6-phosphate, indicating that sugars with carboxyl or phosphate group at their C-6 position are imported by other transport systems distinct from the PTS (24). Despite the identification of more than twenty sugars that are imported by the PTS, none are modified at the C-6 position (25).

The PTS is thought to import depolymerized hyaluronan as the presence of hyaluronan leads to an increase in the expression of the *S. agalactiae* PTS gene (26). Marion *et al*. have previously shown that the PTS, in conjunction with hyaluronate lyase and UGL, is essential for the growth of *S. pneumoniae* when hyaluronan is the sole carbon source (27). PTS mutation has been shown to reduce the ability of the bacteria to colonize mouse upper respiratory tracts. However, the PTS mutant was found to grow on GlcUA and GlcNAc at the same rate as the parental wild-type strain; the substrate of the PTS in this case remains to be identified. This study focused on the role of the *S. agalactiae* PTS in the import of unsaturated hyaluronan disaccharides.

## RESULTS

### Degradation of GAGs by *S. agalactiae*

As *S. agalactiae* produces hyaluronate lyase that depolymerizes both hyaluronan and chondroitin sulfate (28), the halo plate method was used to investigate streptococcal GAG degradation (Fig. 2A). Chondroitin sulfate is classified into chondroitin sulfates A, B, and C, based on the position of the sulfate group (29). Chondroitin sulfate C is sulfated at the C-6 position of GalNAc, whereas chondroitin sulfates A and B are sulfated at the C-4 position. The repeating units of chondroitin sulfates A, B, and C are GlcUA-GalNAc4S (GalNAc with a sulfate group at the C-4 position), IdoUA-GalNAc4S, and GlcUA-GalNAc6S (GalNAc with a sulfate group at the C-6 position), respectively (30). Plates containing the brain heart infusion necessary for streptococcal growth failed to produce the white precipitate that results from the aggregation of GAGs and bovine serum albumin (BSA) upon the addition of acetic acid. Accordingly, nutrient medium and horse serum were used as alternatives for halo plate analysis. In addition to *S. agalactiae* NEM316 (Fig. 1B, upper), *S. agalactiae* JCM 5671 (which contains the GAG genetic cluster; Fig. 1B, lower) was selected to represent a typical strain that is able to degrade GAG. Moreover, the function of the PTS encoded in the GAG genetic cluster was characterized through the construction of a NEM316 mutant strain by replacing the PTS gene segment (a set of EIIB, EIIC, and EIID genes) with a kanamycin-resistant gene (Km^r^) (Fig. 3A); the degrading ability of this PTS mutant was then assessed.

**FIG 2.**
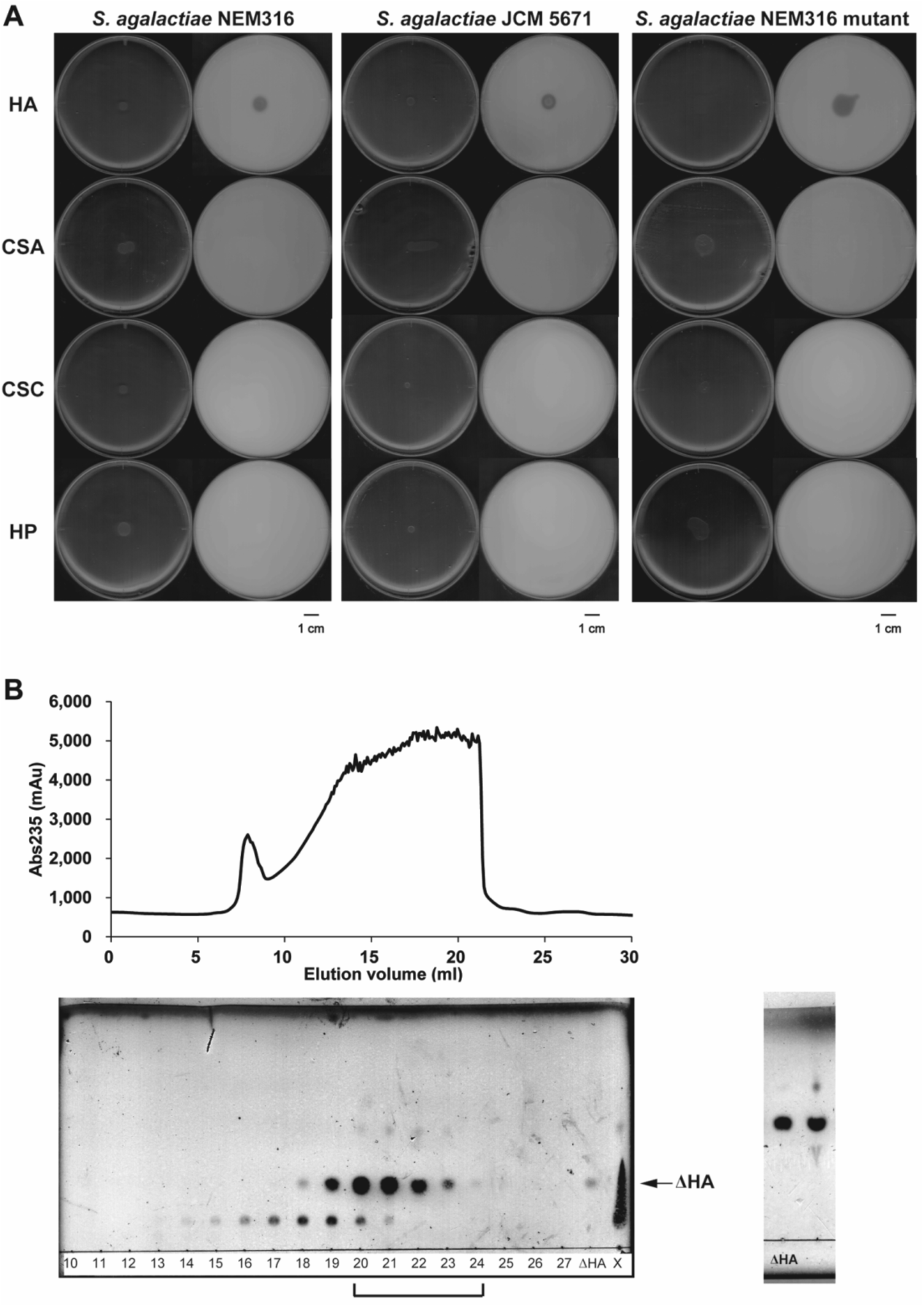
Degradation of GAGs by *S. agalactiae*. (A) Degradation of GAGs by *S. agalactiae* NEM316, *S. agalactiae* JCM 5671, and *S. agalactiae* NEM316 PTS mutant. The left and right plates in each panel are images taken before and after the addition of acetic acid, respectively. Plates contained hyaluronan (HA), chondroitin sulfate A (CSA), chondroitin sulfate C (CSC), or heparin (HP). (B) Preparation of unsaturated hyaluronan disaccharide. Shown are the elution profiles of unsaturated hyaluronan disaccharide during gel filtration chromatography (upper), and TLC profiles of fractions from gel filtration chromatography (lower, left). Numbers denote elution volume (ml). X represents a sample of the reaction mixture before gel filtration chromatography. TLC profiles of the mixture of collected fractions are also shown (lower, right). The mixture (lane, right) was found to contain a saccharide as a main product that corresponded to the standard unsaturated hyaluronan disaccharide (?HA) (lane, left).

**FIG 3.**
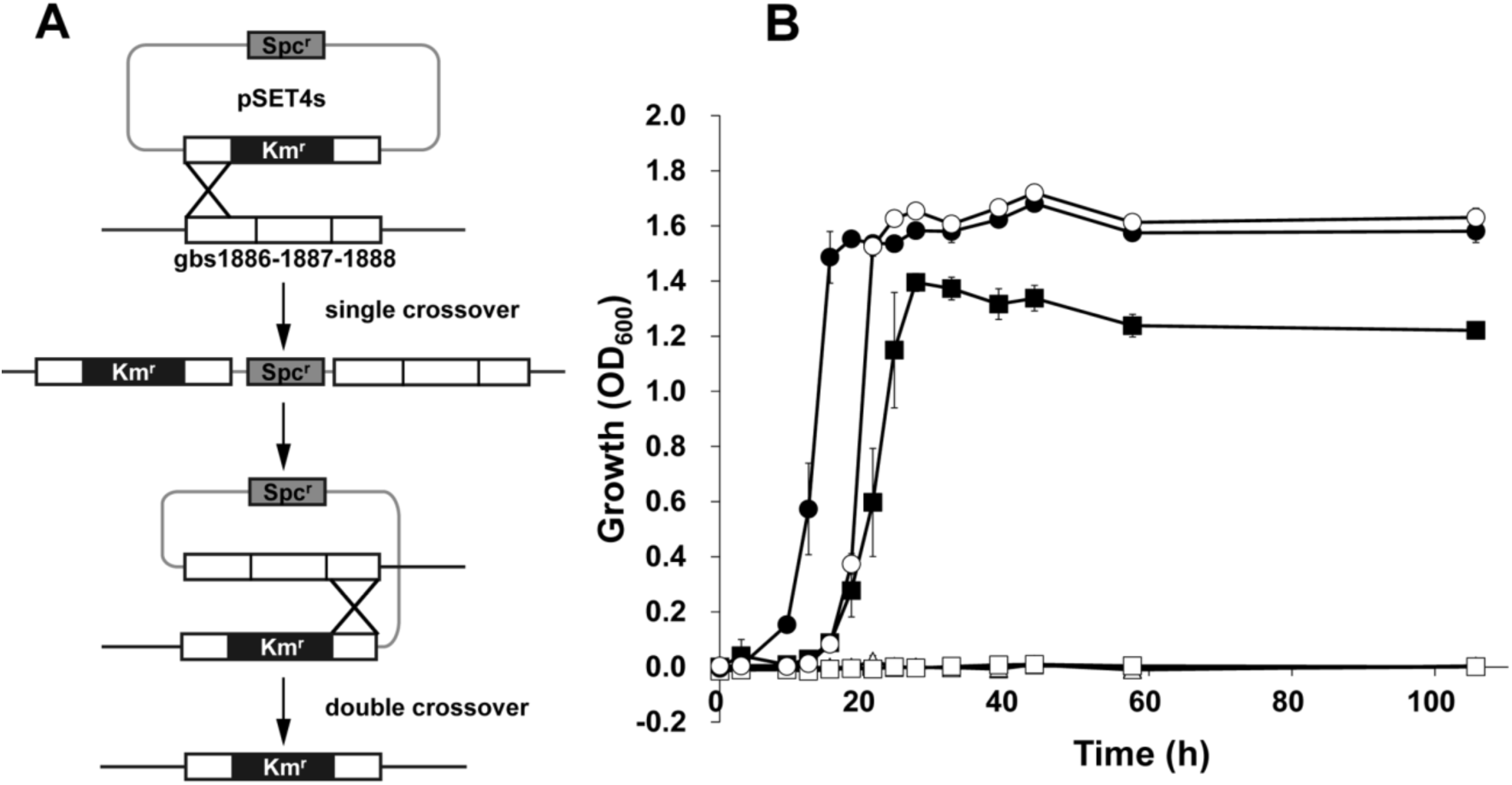
Growth of *S. agalactiae* in the presence of hyaluronan. (A) Construct of the PTS mutant. The gbs1886-1887-1888 operon gene in the pSET4s plasmid was disrupted by the insertion of Km^r^. The pSET4s-gbs1886-1887-1888::Km^r^ plasmid was introduced into the streptococcal cells. A double crossover mutant was obtained by homologous recombination. (B) Wild-type (closed) and the PTS mutant (open) in minimum medium containing hyaluronan (square), glucose (circle), or no saccharide (triangle).

Although the GAG genetic cluster of *S. agalactiae* JCM 5671 is divided into two segments by the insertion of 55 genes between the HMPREF9171_0332 gene encoding the PTS EIIA and the HMPREF9171_0388 gene encoding UGL (Fig. 1B, lower), both strains produced clear halos on plates containing hyaluronan (Fig. 2A). However, halos were not observed on plates containing chondroitin sulfate A or C, or heparin. This indicates that *S. agalactiae* is active against hyaluronan, but not the other three GAGs. The lack of chondroitin sulfate A and C degradation was probably due to a low level of bacterial lyase activity toward chondroitin sulfates. Similar to the wild-type strain, the PTS mutant exhibited a halo on hyaluronan-containing plates (but not on those containing the other GAGs), suggesting that the PTS is not essential for the degradation of hyaluronan.

### Assimilation of hyaluronan by *S. agalactiae*

*S. agalactiae* GD201008-001 has been shown to use hyaluronan as a sole carbon source for growth (31). In addition, a *S. pneumoniae* mutant with a disruption of the PTS genes in the GAG genetic cluster was unable to assimilate hyaluronan (27). Based on these observations, the hyaluronan assimilation of *S. agalactiae* NEM316 (Fig. 1B, upper) and its PTS mutant was investigated using hyaluronan-containing minimum medium (Fig. 3B). *S. agalactiae* was found to grow on hyaluronan or glucose, whereas no growth was apparent on minimum medium that lacked saccharide. In contrast, the PTS mutant was unable to grow in the hyaluronan-containing minimum medium, indicating that the PTS encoded in the GAG genetic cluster is crucial for the assimilation of hyaluronan in *S. agalactiae*. In addition, the growth of wild-type cells on hyaluronan-containing media was higher than that observed in the absence of hyaluronan, whereas the growth of the PTS mutant cells was unaffected (Fig. S2).

As *S. agalactiae* was found to degrade and assimilate hyaluronan, PTS activity was investigated by the preparation of unsaturated hyaluronan disaccharide using recombinant bacterial hyaluronate lyase. An overexpression system for *S. agalactiae* hyaluronate lyase was constructed in *Escherichia coli*, and the cell extract was used to treat hyaluronan. The reaction product was then purified by gel filtration chromatography (Fig. 2B, upper). The eluted fractions were subjected to thin-layer chromatography (TLC) (Fig. 2B, lower), and the fractions containing unsaturated hyaluronan disaccharide at an elution volume of 20–24 ml were collected, concentrated, and used as the substrate in the PTS assay.

### Import of unsaturated hyaluronan disaccharide by *S. agalactiae* PTS

To demonstrate the PTS-dependent import of unsaturated hyaluronan disaccharide in *S. agalactiae* NEM316, PTS-induced pyruvate production from phosphoenolpyruvate was measured using bacterial cells permeabilized by treatment with toluene (Fig. 4). As *S. pneumoniae* has previously been shown to incorporate cellobiose via a PTS (32), cellobiose was used as a positive control. In contrast, D-glucosamine-6-phosphate (GlcN6P) was used as a negative control as phosphorylation at the C-6 position renders it an unsuitable PTS substrate. Gram-positive *Micrococcus luteus*, which lacks both the GAG genetic cluster and PTS genes, was also used as a negative control.

**FIG 4.**
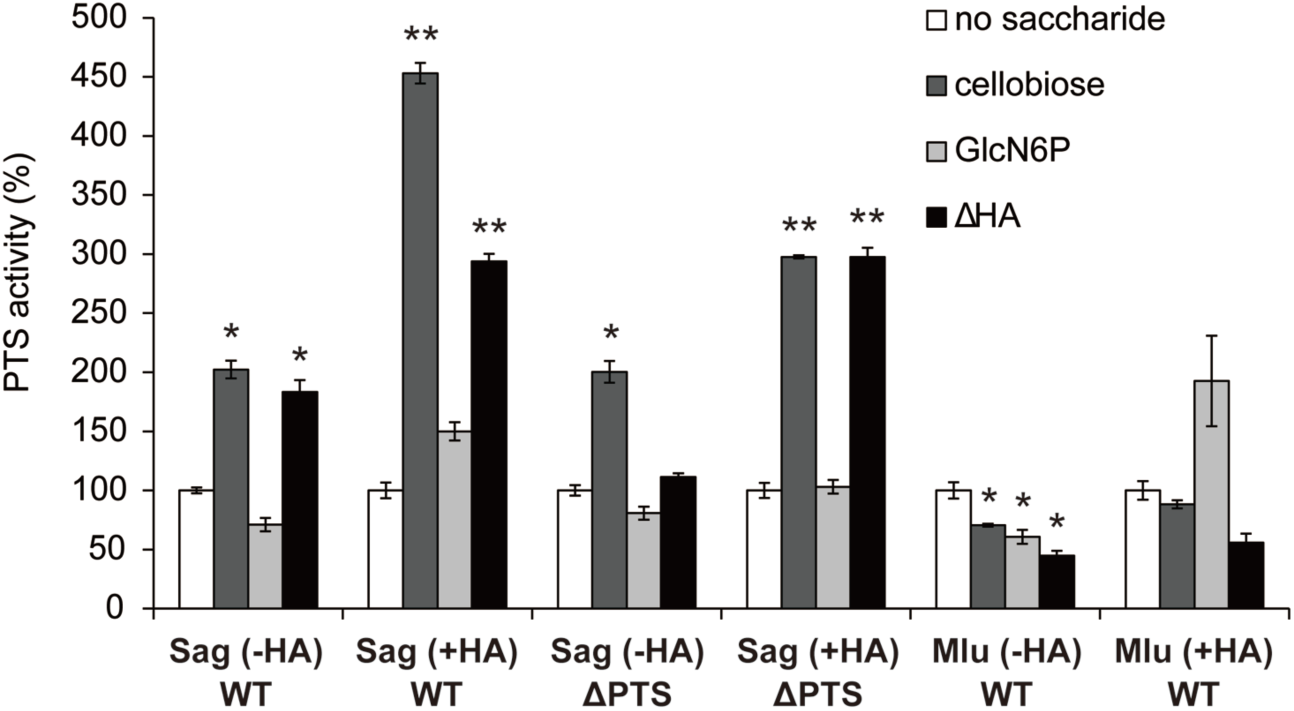
Import of unsaturated hyaluronan disaccharide by *S. agalactiae* PTS. Levels of PTS import into *S. agalactiae* and *M. luteus* grown in the absence or presence of hyaluronan. *S. agalactiae* wild-type cells grown in the absence of hyaluronan, Sag (-HA) WT; *S. agalactiae* wild-type cells grown in the presence of hyaluronan, Sag (+HA) WT; *S. agalactiae* PTS mutant cells grown in the absence of hyaluronan, Sag (-HA) ΔPTS; *S. agalactiae* PTS mutant cells grown in the presence of hyaluronan, Sag (+HA) ΔPTS; *M. luteus* wild-type cells grown in the absence of hyaluronan, Mlu (-HA) WT; and *M. luteus* wild-type grown in the presence of hyaluronan, Mlu (+HA) WT. No saccharide, white; positive control (cellobiose), dark gray; negative control [glucosamine-6-phosphate (GlcN6P)], light gray; and unsaturated hyaluronan disaccharide (ΔHA), black. Each measurement represents the mean of three individual experiments (means ± standard deviations). A significant difference was statistically determined using Student’s t-test (**p < 0.01; *p < 0.05).

The growth of *S. agalactiae* both in the presence and absence of hyaluronan led to an increase in the PTS import of cellobiose (compared to the basal activity measured in the absence of the sugar substrate). This indicates that the bacterial PTS is promoting the uptake of cellobiose into the cell. In contrast, *S. agalactiae* exhibited no enhanced PTS activity using GlcN6P as a substrate, regardless of the presence of hyaluronan. These results suggest that permeabilized *S. agalactiae* cells are functionally active, and the assay used is a reliable indicator of PTS activity. The bacterial cells exhibited a higher level of PTS-mediated cellobiose import when grown in the presence of hyaluronan; this is probably due to hyaluronan-dependent increases in the transcriptional level of the PTS genes for import of cellobiose (26). In contrast, *M. luteus* exhibited similar levels of PTS import of cellobiose as the basal controls; this reflects the lack of a cellobiose PTS in *M. luteus*. Limited PTS import of GlcN6P was observed in *M. luteus* cells, which is in agreement with the results from *S. agalactiae*.

The levels of the PTS import of unsaturated hyaluronan disaccharide into *S. agalactiae* grown in the absence and presence of hyaluronan were approximately 1.8 and 2.9 times higher than basal levels, respectively. These findings represent a significant difference between the PTS import and basal activity, especially in cells grown in the presence of hyaluronan. On the other hand, the PTS mutant grown in the absence of hyaluronan showed similar levels of PTS import of unsaturated hyaluronan disaccharide as the basal controls. This indicates that the mutant lacks the ability to import unsaturated hyaluronan disaccharide. These results clearly demonstrate that *S. agalactiae* imports unsaturated hyaluronan disaccharide using the PTS encoded in the GAG genetic cluster.

### Structure determination of *S. agalactiae* EIIA

X-ray crystallography of EIIA^ΔHA^ (EIIA for unsaturated hyaluronan disaccharide; ΔHA) was performed as a first step toward determining the overall structure of the *S. agalactiae* PTS complex (EIIABCD) for the import of unsaturated hyaluronan disaccharide. Recombinant purified EIIA^ΔHA^ protein was crystallized, and X-ray diffraction data were collected. Data collection and refinement statistics are shown in Table 1. The EIIA^ΔHA^ crystal belongs to the *P*1 group with unit cell dimensions of *a* = 52.3, *b* = 53.8, and *c* = 94.9 Å, and *α* = 91.1, *β* = 90.0, and *γ* = 61.0°. The final model, containing six molecules in an asymmetric unit, was refined to an *R*_work_ of 20.8% up to a resolution of 1.8 Å. Ramachandran plot analysis indicated 99.0% of residues in the favored regions and 1.00% of residues in the additional allowed regions. The crystal structure of EIIA^ΔHA^ was determined using molecular replacement with *E. coli* PTS EIIA^man^ (PDB ID, 1PDO) for mannose import as the initial model.

**TABLE 1.**
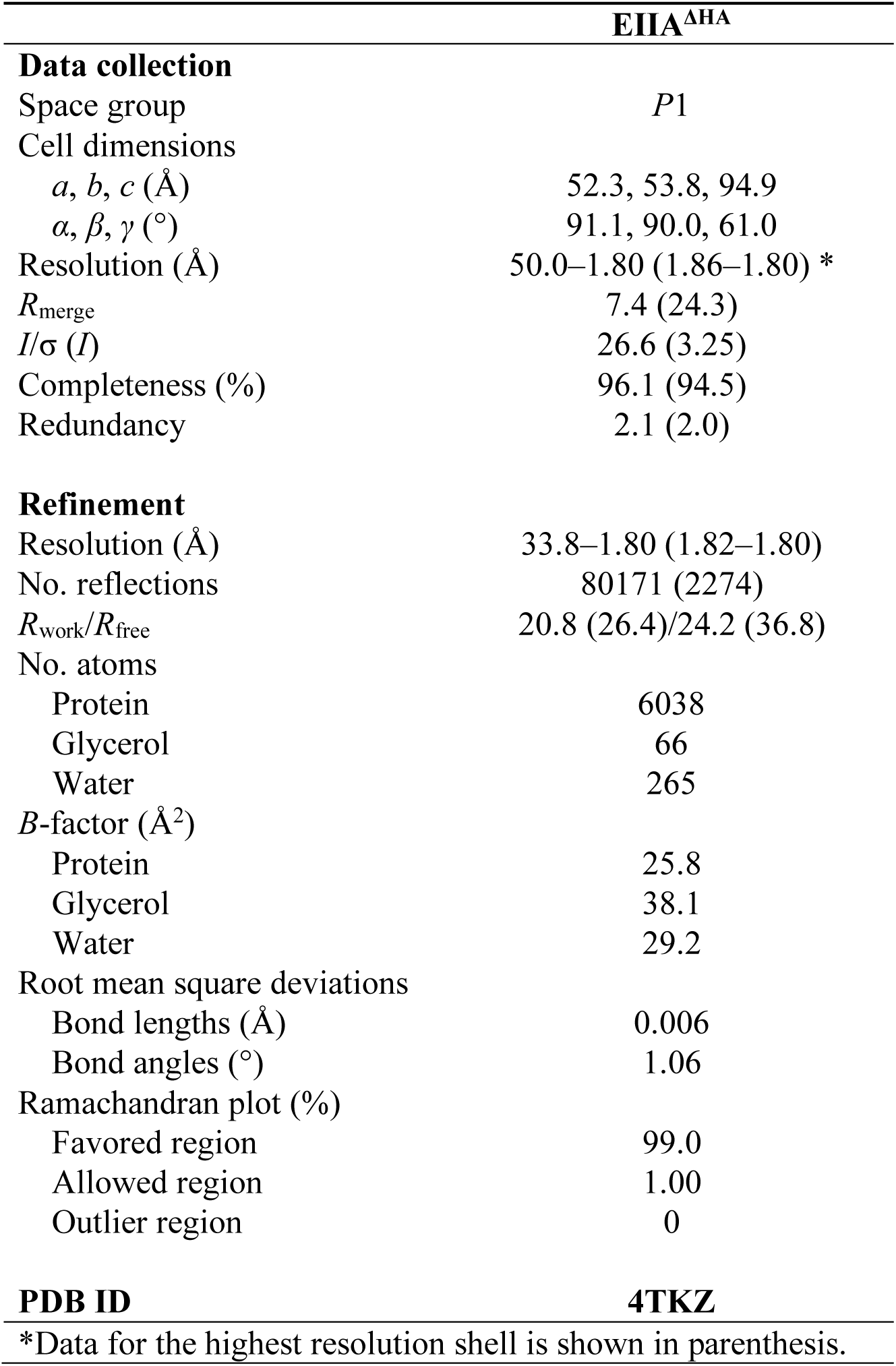
Statistics of EIIA^ΔHA^ for data collection and structure refinement

### Crystal structure of EIIA^**ΔHA**^

EIIA^ΔHA^ consists of 144 residues, although the 14 residues (Leu131–Ile144) of the C-terminal could not be assigned due to their structural flexibility. With respect to the secondary structure of EIIA^ΔHA^, α-helices, β-sheets, and loops constitute 41.0%, 17.0%, and 42.0%, respectively. EIIA^ΔHA^ is composed of six α-helices (α1, Phe12–Ala24; α2, Ser41–Val52; α3, Thr68–Leu76; α4, Leu93–Met105; α5, Asp110–Glu122; and α6, Phe127– Thr129), five β-strands (β1, Lys3–His9; β2, Val30–Phe35; β3, Glu57–Thr62; β4, Lys84–Ser89; and β5, Val125–Asp126), and ten loops (L1, Met1–Ile2; L2, Gly10–Asn11; L3, Gly25–Tyr29; L4, Ile36–Ser40; Ile53–Lys56; L5, Asp63–Gly67; L6, Ser77–Lys83; L7, Gly90–Asn92; L8, Phe106–Val109; L9, Gly123–Ile124; and L10, Cys130). In the overall structure, a parallel β-sheet containing four β-strands (β1, β2, β3, and β4) is located at the center, and two (α2 and α3) and three α-helices (α1, α4, and α5) are located so they pinch the β-sheet from both sides, resulting in the formation of a Rossman-fold frame (Fig. 5A). β1–4 and α1–4 are alternately arranged and α4 is followed by α5 then β5. Gel filtration chromatography suggested that EIIA^ΔHA^ was smaller than a tetramer, and the biological asymmetric unit was shown to be a dimer using *PISA* software (33). In the dimer, the C-terminal β5s in adjacent monomers are arranged to align with mutual β4s, and added to the parallel β-sheet located at the center of the monomer, forming an antiparallel β-sheet.

**FIG 5.**
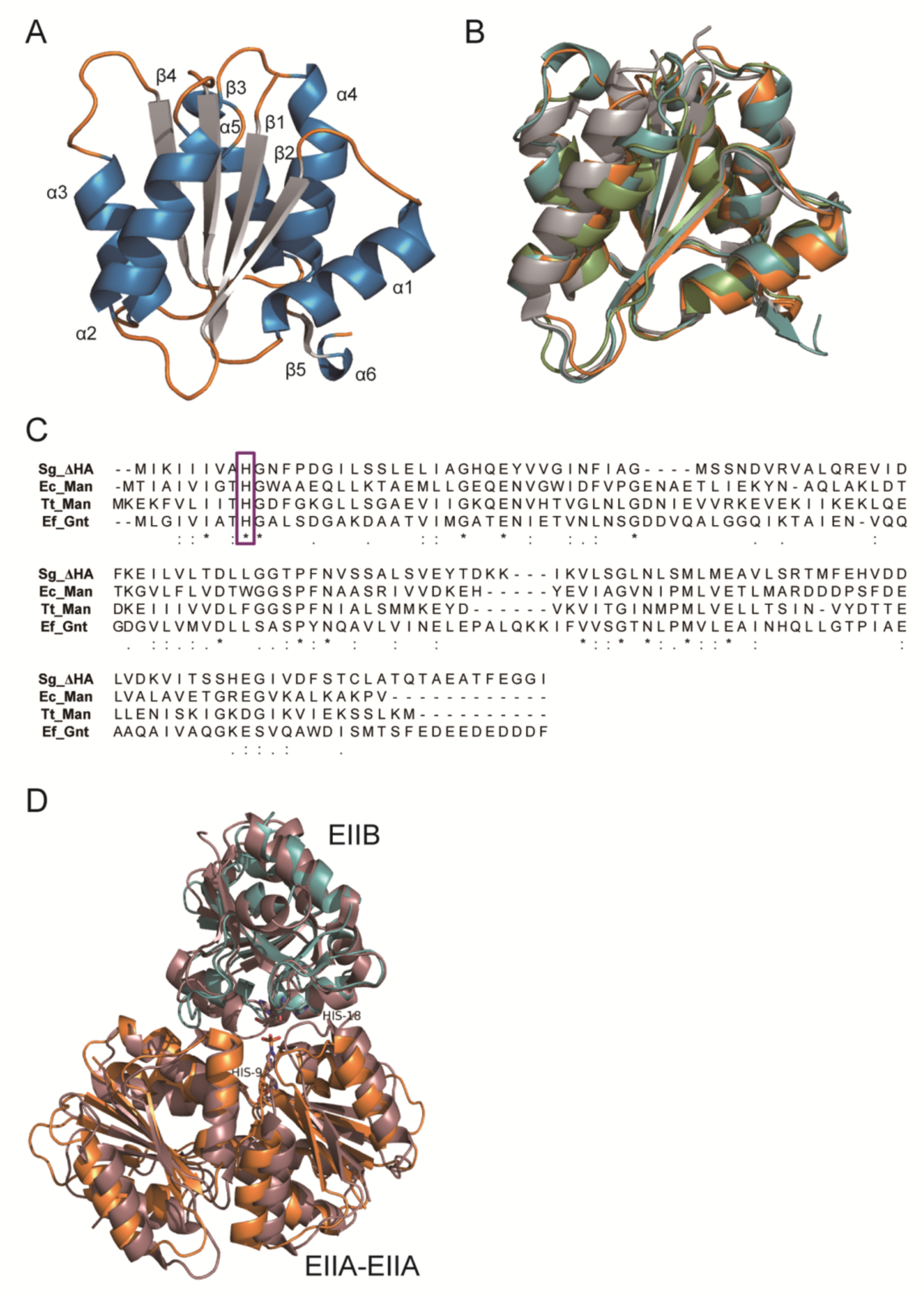
Three-dimensional structure of *S. agalactiae* PTS EIIA^ΔHA^. (A) Overall structure of *S. agalactiae* EIIA^ΔHA^. Blue, α-helices; gray, β-strands; and orange, loops. (B) Homologue proteins of *S. agalactiae* EIIA^ΔHA^. Orange, *S. agalactiae* EIIA^ΔHA^ (Sg_ΔHA); light green, *E. coli* mannose EIIA^Man^ (Ec_Man); gray, *T.tengcongensis* mannose EIIA^Man^ (Tt_Man); and cyan, *E. faecalis* gluconate EIIA^Gnt^ (Ef_Gnt). (C) Primary structure alignment of EIIA. (D) Superimposition of *S. agalactiae* EIIA^ΔHA^-EIIB^ΔHA^ and *E. coli* EIIA^Man^-EIIB^Man^ complexes. Orange, *S. agalactiae* EIIA^ΔHA^; cyan, *S. agalactiae* EIIB^ΔHA^ (modeling); dark pink, *E. coli* EIIA^Man^-EIIB^Man^ complex.

## DISCUSSION

The ability of *S. agalactiae* to degrade and assimilate hyaluronan allows the measurement of PTS import activity using an unsaturated disaccharide derived from hyaluronan degradation. The PTS import of unsaturated hyaluronan disaccharide in bacterial wild-type cells grown in the absence of hyaluronan was significantly higher than in controls using no substrate or GlcN6P; this indicates that *S. agalactiae* incorporates unsaturated hyaluronan disaccharide via a PTS. On the other hand, PTS mutant cells grown in the absence of hyaluronan were unable to incorporate unsaturated hyaluronan disaccharide. Based on these observations and the fact that PTS mutant failed to grow on hyaluronan-containing minimum medium, we conclude that the PTS encoded in the GAG genetic cluster is the sole importer of unsaturated hyaluronan disaccharide in *S. agalactiae*.

Surprisingly, the PTS mutant grown in the presence of hyaluronan exhibited enhanced PTS import of unsaturated hyaluronan disaccharide, in comparison with the control. Furthermore, the levels of PTS import of unsaturated hyaluronan disaccharide and cellobiose in bacterial wild-type cells grown in the presence of hyaluronan, were approximately 1.6 and 2.3 times higher than those in the absence of hyaluronan, respectively. Based on these observations, we hypothesize that hyaluronan present in the cultured medium damages growing *S. agalactiae* cells, and the subsequent toluene-treatment causes leakage of cytoplasmic enzymes such as UGL and β-glucosidase. As a result, the unsaturated hyaluronan disaccharide and cellobiose contained in the reaction mixtures are degraded by these enzymes to the constituent monosaccharides (unsaturated GlcUA, GlcNAc, and glucose), and incorporated by another PTS.

While bacterial cells import sugars through various mechanisms such as facilitated diffusion, primary and secondary active transport, and group translocation, the PTS is the major sugar import pathway in many species (34, 35). PTS Enzyme II is classified into four families based on its primary structure: (i) the glucose-fructose-lactose family; (ii) the ascorbate-galactitol family; (iii) the mannose family; and (iv) the dihydroxyacetone family (36). Several characteristic features of the mannose family have been defined. These include the observations that EIIC is a hetero (not a homo)-membrane domain in combination with EIID, an EIIB receives a phosphate group from a histidine rather than a cysteine residue, and various sugars can be used as a substrate. *S. agalactiae* EIIA^ΔHA^ showed the most similarity with *E. coli* mannose EIIA (PDB ID, 1PDO), *Enterococcus faecalis* gluconate EIIA (PDB ID, 3IPR), and *Thermoanaerobacter tengcongensis* mannose/fructose EIIA (PDB ID, 3LFH), all of which belong to the mannose family; Z-scores, estimated by the Dali program (37), were 20.2, 19.9, and 19.1, respectively (Figs. 5B and C) (Table S1). Based on the well-conserved characteristics of the mannose family, the *S. agalactiae* PTS for the import of unsaturated hyaluronan disaccharide appears to be a member of this family. The three-dimensional structures of these enzymes were well superimposed (Fig. 5B). *S. agalactiae* EIIB^ΔHA^ was homology modeled by the SWISS-MODEL using putative *S. pyogenes* EIIB for the import of GalNAc (PDB ID, 3P3V) (sequence identity: 70%) as a template (38). *S. agalactiae* EIIB^ΔHA^ is composed of an antiparallel β-sheet of eight β-strands, and eight α-helices (Fig. 5D). The three-dimensional structures of *E. coli* HPr-EIIA^Man^ and EIIA^Man^-EIIB^Man^ complexes (previously determined using NMR), and EIIA^Man^, have also been found to form a dimer (39, 40). In HPr-EIIA^Man^ and EIIA^Man^-EIIB^Man^ complexes, His10 of EIIA^Man^ is an important residue in the transfer of a phosphate group from HPr to EIIB^Man^, through EIIA^Man^ (41). Due to the similarity between the interaction sites of EIIA^Man^ with HPr, and EIIA^Man^ with EIIB^Man^, HPr or EIIB^Man^ must be separated while the other remains bound to EIIA^Man^. The His10 of EIIA^Man^ is also conserved in EIIA^ΔHA^ with His9 (Fig. 5C). To compare EIIA^ΔHA^-EIIB^ΔHA^ and EIIA^Man^-EIIB^Man^, the EIIA^ΔHA^ dimer and EIIB^ΔHA^ monomer were superimposed with the EIIA^Man^-EIIB^Man^ complex. The arrangements of the His9 of EIIA^ΔHA^ and His10 of EIIA^Man^ almost corresponded to each other; this was also observed with the His18 of EIIB^ΔHA^ and His18 of EIIB^Man^. His9 of EIIA^ΔHA^ is located at the end of β1 and interacts with Asp63 via hydrogen bonding, and with Phe12, Phe35, Asp63, Gly67, and Pro69 through van der Waals contacts. These amino acid residues are almost conserved in EIIA for mannose (including *E. coli* EIIA^Man^). Therefore, His9 of EIIA^ΔHA^ appears to be crucial in the transfer of a phosphate group.

In this study, *S. agalactiae* was found to import unsaturated hyaluronan disaccharide through the PTS encoded in the GAG genetic cluster. Unlike other sulfated GAGs, hyaluronan contains no sulfate groups at the C-6 position of its constituent monosaccharides. Thus, unsaturated hyaluronan disaccharide is a suitable substrate for the PTS through the transfer of a phosphate group to the C-6 position. We have recently identified a solute-binding protein-dependent ABC transporter in Gram-negative *Streptobacillus moniliformis* that acts as an importer of unsaturated GAG disaccharides (42, 43) (Fig. S3). Bacterial ABC transporters generally receive substrates from solute-binding proteins and incorporate the substrates into the cytoplasm using the energy of ATP hydrolysis (44). Because the imported substrates of the ABC transporter have no modifications that render them distinct from PTS substrates, *S. moniliformis* ABC transporter has been demonstrated to import both sulfated and non-sulfated unsaturated GAG disaccharides derived from chondroitin sulfate and hyaluronan. Furthermore, genes homologous with *S. moniliformis* ABC transporter genes are conserved among the genome of several fusobacterial species, which are generally indigenous to animal oral cavities. Fusobacterium probably utilizes the ABC transporter for the assimilation of sulfated GAGs that are abundant in the oral cavities. On the other hand, *S. agalactiae,* a pathogen of the hyaluronan-rich vaginal mucosa (45, 46), is thought to utilize the PTS to assimilate vaginal hyaluronan. Streptococci, and several intestinal probiotics such as *Lactobacillus rhamnosus* and *E. faecalis*, possess the genes homologous with the PTS (47), indicating that the import system may be common to the intestinal bacteria that are able to use GAGs.

In conclusion, this is the first report that confirms *S. agalactiae* PTS encoded in the GAG genetic cluster is the importer of non-sulfated unsaturated hyaluronan disaccharides distinct from sulfated GAG-fragments.

## MATERIALS AND METHODS

### Materials

Hyaluronan sodium salt was purchased from Sigma Aldrich. Sodium salts of chondroitin sulfates A and C were obtained from Wako Pure Chemical Industries, and heparin sodium salt from Nacalai Tesque. A thermosensitive suicide vector, pSET4s, was kindly provided by Dr. Takamatsu (National Agriculture and Food Research Organization).

### Micoorganisms and culture conditions

*S. agalactiae* NEM316 (ATCC 12403) was purchased from Institute Pasteur, and *S. agalactiae* JCM 5671 (ATCC 13813) from Riken BioResource Center. *S. agalactiae* cells were statically grown at 37°C under 5% CO_2_ in 3.7% brain heart infusion medium (BD Bacto) or 0.8% nutrient medium (0.3% beef extract and 0.5% peptone) (Difco), supplemented with 20% horse serum for 16–24 h. To investigate hyaluronan assimilation by *S. agalactiae*, hyaluronan-containing minimum medium was prepared as described previously (48). Briefly, streptococcal cells in logarithmic phase of growth were inoculated (to an optical density of 0.01 at 600 nm; OD_600_) into minimum medium consisting of 0.44 g/l KH_2_PO_4_, 0.3 g/l K_2_HPO_4_, 3.15 g/l Na_2_HPO_4_, 2.05 g/l NaH_2_PO_4_, 0.225 g/l sodium citrate, 6 g/l sodium acetate, 0.6 g/l (NH_4_)_2_SO_4_, 0.2 g/l MgSO_4_, 10 mg/l NaCl, 10 mg/l FeSO_4_, 10 mg/l MnSO_4_, 0.4 mg/l riboflavin, 0.01 mg/l biotin, 0.1 mg/l folate, 0.8 mg/l pantothenate, 0.4 mg/l thiamine, 2 mg/l nicotinamide, 0.8 mg/l pyridoxamine, 0.1 mg/l *p*-aminobenzoate, 5 mg/l Gln, 300 mg/l Glu, 110 mg/l Lys, 100 mg/l Asp, 100 mg/l Ile, 100 mg/l Leu, 100 mg/l Met, 100 mg/l Ser, 100 mg/l Phe, 100 mg/l Thr, 100 mg/l Val, 200 mg/l Ala, 200 mg/l Arg, 200 mg/l Cys, 200 mg/l His, 200 mg/l Gly, 400 mg/l Pro, 200 mg/l Trp, 200 mg/l Tyr, 35 mg/l adenine and 30 mg/l uracil, in the presence of 0.5% hyaluronan or 2% glucose, or in the absence of sugar substrate. The bacterial cells were grown at 37°C and turbidity monitored periodically.

*E. coli* DH5α harboring plasmids were cultured at 37°C in Luria-Bertani (LB) medium containing 100 µg/ml sodium ampicillin. For the expression of recombinant proteins, *E. coli* BL21(DE3) harboring plasmids were cultured at 30°C in LB medium containing 100 µg/ml sodium ampicillin to an OD_600_ of 0.3–0.7, followed by the addition of isopropyl-β-D-thiogalactopyranoside to a final concentration of 0.1 mM, and further incubation at 16°C for 2 days.

### Halo detection for GAG degradation

The halo detection method was used to investigate the GAG-degrading ability of *S. agalactiae*. The bacterial cells were grown on plates containing 0.2% dialyzed GAG (hyaluronan, chondroitin sulfate A, C, or heparin), 0.8% nutrient medium, 20% horse serum, and 1% BSA solidified with 1% agar. When sufficient bacterial growth was achieved, the addition of 2 M acetic acid (1 ml) to the plates resulted in the formation of a white precipitate due to the interaction of GAGs and BSA; areas containing degraded GAGs appear as clear zones or “halos.”

### Construction of an overexpression system

An overexpression system for *S. agalactiae* hyaluronate lyase was constructed in *E. coli* as a source of enzymes for the preparation of unsaturated hyaluronan disaccharide required for the PTS import assay. To clone the gbs1270 gene that encodes hyaluronate lyase, polymerase chain reaction (PCR) was conducted on 10 µl of reaction mixture consisting of 0.2 U of KOD Plus Neo polymerase (Toyobo), *S. agalactiae* as a template, 0.3 pmol of each of forward and reverse primers, 2 nmol of dNTPs, 10 nmol of MgCl_2_, 0.5 µl of dimethyl sulfoxide, and the commercial reaction buffer supplied with KOD Plus Neo polymerase. PCR conditions were as follows: 94°C for 2 min followed by 30 cycles of 98°C for 10 s, 35°C for 30 s, and 68°C for 2 min. The PCR product was ligated to *Hin*cII-digested pUC119 (Takara Bio) using Ligation High Ver. 2 (Toyobo), and the resulting plasmid was digested with *Nco*I and *Xho*I to isolate the gbs1270 gene. The gene fragment was confirmed to encode the correct gbs1270 by DNA sequencing (49). The *Nco*I and *Xho*I-digested gbs1270 gene was ligated into *Nco*I and *Xho*I-digested pET21d (Novagen), and *E. coli* BL21(DE3) host cells were transformed with the resulting plasmid, pET21d-gbs1270. An overexpression system of *S. agalactiae* EIIA^ΔHA^ was also constructed in *E. coli* for X-ray crystallography. To clone the gbs1890 gene encoding EIIA^ΔHA^, PCR was performed using primers specific for EIIA^ΔHA^ as described above. The gene fragment was ligated into *Nde*I and *Xho*I-digested pET21b (Novagen), and *E. coli* BL21(DE3) host cells were transformed with the plasmid pET21b-gbs1890. The PCR primers used are shown in Table 2.

**TABLE 2.**
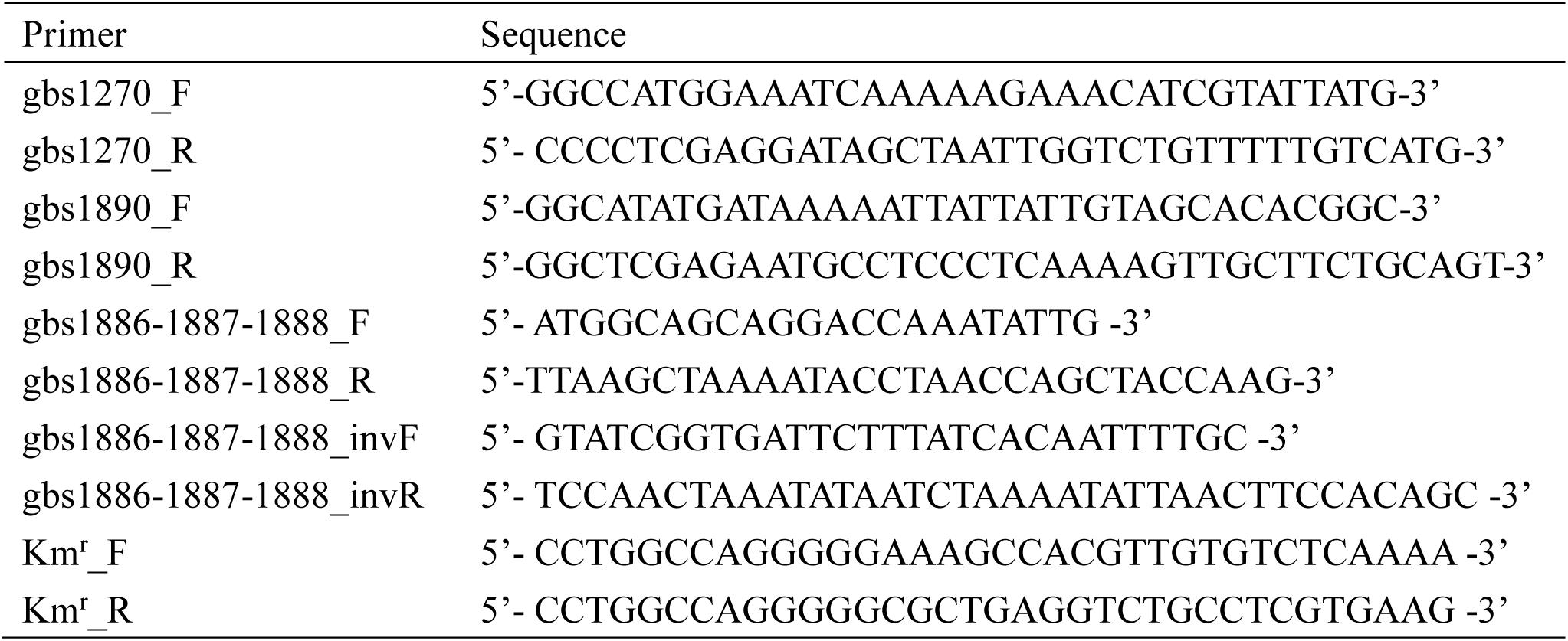
Primers used in this study Primer

### Construction of the PTS mutant

The PTS mutant was constructed using Km^r^ and the thermosensitive suicide vector, pSET4s. The gbs1886-1887-1888 operon gene coding for PTS EIID, EIIC, and EIIB was amplified by PCR using *S. agalactiae* as a template, and the PCR product was ligated with *Hin*cII-digested pSET4s. Using the pSET4s-gbs1886-1887-1888 plasmid as a template, inverse PCR was conducted to amplify lineared PCR product to remove the PTS operon gene excepting the both ends of 500 bp for homologous recombination. The lineared product of inverse PCR was ligated with pUC4K-derived Km^r^. The resulting plasmid was designed as pSET4s-gbs1886-1887-1888::Km^r^.

To transform the pSET4s-gbs1886-1887-1888::Km^r^ plasmid into the streptococcal cells, electrotransformation was conducted as previously described, but with a slight modification (50). Briefly, *S. agalactiae* NEM316 was grown in 50 ml brain heart infusion medium containing 0.4% glucose to an OD_600_ of 0.3, harvested by centrifugation at 2,610 *g* at 4°C for 10 min, and washed three times with 15 ml of 10% cold glycerol. The washed cells were suspended in 1 ml of 20% cold glycerol and aliquoted into 50 µl samples. Following the addition of 1 µg of the plasmid (1 µg/µl) and incubation on ice for 1 min, the competent cells were transferred to a cold electroporation cuvette with a 0.1 cm gap (Bio-Rad). The cuvette was set to MicroPulser (Bio-Rad) and pulsed as follows: field strength, 1.8 kV; capacitor, 10 µF; and resistor, 600 Ω. Brain heart infusion medium containing 10% glycerol (1 ml) was immediately and gently added to the cuvette. After incubation at 28°C for 1 h, the electroporated cells were spread on a brain heart infusion plate containing 250 µg/ml spectinomycin, and further incubated at 28°C for 3 d to obtain a spectinomycin-resistant transformant. The single crossover mutant cells obtained were transferred to medium without spectinomycin and subcultured repeatedly at 37°C. A spectinomycin-sensitive and kanamycin-resistant (500 µg/ml) single colony was considered to be double crossover mutant.

### Protein purification

Recombinant *E. coli* cells were harvested by centrifugation at 6,700 *g* at 4°C for 10 min and suspended in 20 mM Tris (hydroxymethyl) aminomethane-hydrochloride (Tris-HCl), pH 7.5. The cell suspension was ultrasonicated (Insonator Model 201M, Kubota) at 0°C and 9 kHz for 10 min, and subjected to centrifugation at 20,000 *g* at 4°C for 20 min. The supernatant cell extract was then used in subsequent experiments. The BL21(DE3)/pET21d-gbs1270 cell extract was used as a source of hyaluronate lyase for the preparation of unsaturated hyaluronan disaccharide. EIIA^ΔHA^ was purified from BL21(DE3)/pET21b-gbs1890 cell extract using metal affinity [TALON (Clontech)] and gel filtration chromatography [Sephacryl S-200 (GE Healthcare)]. After the confirmation of protein purity by sodium dodecyl sulfate-polyacrylamide gel electrophoresis (SDS-PAGE), the purified protein was dialyzed against 20 mM Tris-HCl (pH 7.5).

### Preparation of unsaturated hyaluronan disaccharide

To investigate PTS import activity, unsaturated hyaluronan disaccharide was prepared using recombinant hyaluronate lyase. A reaction mixture containing BL21(DE3)/pET21d-gbs1270 cell extract, 0.2% hyaluronan, and 20 mM Tris-HCl (pH 7.5) was incubated at 30°C for 2 days. The mixture was then boiled to stop the reaction and centrifuged at 20,000 *g* for 20 min to remove aggregated proteins. The resulting supernatant was concentrated by freeze-drying and subjected to gel filtration chromatography [Superdex Peptide 10/300 GL (GE Healthcare)]. The eluted fractions containing unsaturated hyaluronan disaccharide were identified by monitoring the absorbance (235 nm) from the C=C double bonds of the disaccharide. To confirm the presence of unsaturated hyaluronan disaccharide, pooled fractions were subjected to TLC using a solvent system of 1-butanol:acetic acid:water (3:2:2, v:v:v), and hyaluronan breakdown products were visualized by heating the TLC plates [silica gel 60 F_254_ (Merck)] at 130°C for 5 min after spraying with ethanol containing 10% sulfuric acid. The final disaccharide preparation was freeze-dried and dissolved in sterilized water to a final concentration of 200 mM.

### PTS assay

The import of unsaturated hyaluronan disaccharides into *S. agalactiae* via the PTS was evaluated by quantifying the pyruvate produced from phosphoenolpyruvate during the import process, as described previously but with a few modifications (34, 51). The reaction mixture contained cells whose cell-surface layer had been permeabilized with toluene, phosphoenolpyruvate, disaccharides, NADH, and L-lactate dehydrogenase from rabbit muscle (Oriental Yeast). The reaction was monitored through measurements of absorbance at 340 nm to determine levels of NADH oxidation resulting from the production of lactate from the pyruvate generated by the PTS process. Briefly, *S. agalactiae* wild-type and PTS mutant cells were grown at 37°C under 5% CO_2_ for 16 h in 0.8% nutrient medium and 20% horse serum, in the presence or absence of 0.2% dialyzed hyaluronan, and then harvested by centrifugation at 2,610 *g* at 4°C for 5 min. The cells were washed twice with 5 mM MgCl_2_ and 0.1 M potassium phosphate buffer (KPB); pH 7.2, and suspended in 1 ml of the same buffer containing 50 µl of acetone:toluene (9:1, v:v) by vortexing twice for 2 min. The reaction mixture containing toluene-treated cells, 10 mM sugar, 0.1 mM NADH, 0.023 mg/ml L-lactate dehydrogenase, 10 mM NaF, 5 mM MgCl_2_, and 0.1 M KPB (pH 7.5) was incubated at 37°C for 5 min, and phosphoenolpyruvate then added to a concentration of 5 mM. The reaction was monitored by measuring the decrease in absorbance at 340 nm. The protein concentration of toluene-treated cells was determined using the bicinchoninic acid (BCA) assay (52). The PTS import activity value was calculated as the amount of pyruvate produced (nmol/min/mg).

### X-ray crystallography

To determine the three-dimensional structure of *S. agalactiae* EIIA^ΔHA^, the purified protein was concentrated to 9.24 mg/ml and crystallized using the sitting drop vapor diffusion method. The purified EIIA^ΔHA^ (1 µl) was mixed with an equal volume of a reservoir solution consisting of 20% (w/v) polyethylene glycol (PEG) 3,350 and 0.2 M sodium thiocyanate (pH 6.9), and incubated at 20°C. The crystal was picked up from the drop with a nylon loop, soaked in reservoir solution containing 20% glycerol as a cryoprotectant, and instantaneously frozen in liquid nitrogen. X-ray diffraction data were collected at the BL38B1 station of SPring-8 (Hyogo, Japan). The data were indexed, integrated, and scaled using *HKL2000* software (53). The structure was determined through the molecular replacement method using *Molrep* in the *CCP4* software package and *E. coli* PTS EIIA (PDB ID, 1PDO) for mannose import (54). Structure refinement was conducted with *phenix refine* in *PHENIX* software (55). The model was refined manually with *winCoot* software (56). Protein structures were prepared using the *PyMOL* (57).

## ACKNOWLEDGMENTS

We thank Dr. Daisuke Takamatsu, National Agriculture and Food Research Organization, for kindly supplying a thermosensitive suicide vector, pSET4s. We thank Ms. Ai Matsunami for her excellent technical assistance. We thank Drs. S. Baba and N. Mizuno of the Japan Synchrotron Radiation Research Institute (JASRI) for their kind help in data collection. Diffraction data were collected at the BL38B1 line of SPring-8 (Hyogo, Japan) with the approval of (Projects 2012A1317 and 2016B2574). This work was supported in part by Grants-in-Aid Scientific Research from the Japan Society for the Promotion of Science (to W. H.), and the Targeted Proteins Research Program (to W. H.) from the Ministry of Education, Culture, Sports, Science, and Technology (MEXT) of Japan, and Research Grants (to W. H.) from Yakult Bio-science Foundation. The authors would like to thank Enago (www.enago.com) for the English language review.

## FOOTNOTES

Supplemental material for this article may be found at xxxxx.

